# The Role of Potassium and Host Calcium Signaling in *Toxoplasma gondii* egress

**DOI:** 10.1101/2020.03.06.980508

**Authors:** Stephen A. Vella, Christina A. Moore, Zhu-Hong Li, Miryam A. Hortua Triana, Evgeniy Potapenko, Silvia N J Moreno

## Abstract

*Toxoplasma gondii*, an obligate intracellular parasite, is capable of invading virtually any nucleated cell. Ca^2+^ signaling is universal and both *T. gondii* and its mammalian host cell will utilize Ca^2+^ signaling to stimulate diverse cellular functions. Egress of *T. gondii* from the host cell is an essential step for the infection cycle of *T. gondii* and a cytosolic Ca^2+^ increase initiates the Ca^2+^ signaling cascade that culminates in stimulation of motility and egress. In this work we demonstrate that intracellular *T. gondii* is capable of taking up Ca^2+^ from the host cytoplasm when this concentration is increased during host signaling events. Both intracellular and extracellular Ca^2+^ sources are important to reach a threshold of cytosolic Ca^2+^ needed for a successful egress. Two peaks of Ca^2+^ were observed in single parasites that egressed with the second peak resulting from Ca^2+^ influx. We patched infected host cells to allow a precise delivery of exact concentrations of Ca^2+^ for stimulating motility and egress. Using this approach, we found that low potassium concentration modulates but do not trigger host cell egress. This is the first study using whole-cell patches to study the role of ions such as K^+^ and Ca^2+^ in *T. gondii* egress.

## Introduction

The pathogenesis caused by the infection with *Toxoplasma gondii* is linked to the ability of the parasite to engage in multiple rounds of a lytic cycle which consists of invasion of host cells, replication inside a parasitophorous vacuole (PV), and egress resulting in lysis of the host cell, followed by invasion of a new host cell (1, 2). Several key steps of the lytic cycle of *T. gondii:* egress, invasion, attachment, and motility, are regulated by fluctuations in cytosolic Ca^2+^ concentrations (3, 4).

Ca^2+^ is a universal signaling molecule and plays important roles in the regulation of a large number of cellular functions (5). The cytosolic concentration of Ca^2+^ is highly regulated, because prolonged high cytosolic Ca^2+^ levels are toxic and may result in cell death. A variety of Ca^2+^ pumps, channels, and transporters, located at the plasma membrane (PM) and to intracellular stores (endoplasmic reticulum (ER), acidic stores and mitochondria) are involved in maintaining cytosolic Ca^2+^ levels under control (5).

In *Toxoplasma* Ca^2+^ signaling initiates a chain of events that leads to the activation of specific effectors involved in the regulation of motility as gliding parasites loaded with fluorescent Ca^2+^ indicators, as well as expressing Genetically Encoded Calcium Indicators (GECIs) show Ca^2+^ oscillations (6, 7). Previous studies have shown that increasing cytoplasmic Ca^2+^ activates the motility machinery leading to egress. Blocking these cytosolic Ca^2+^ fluxes using BAPTA-AM (membrane permeable cytosolic Ca^2+^ chelator), blocks motility, conoid extrusion (apical tip of cell necessary for attachment), invasion, and host cell egress (8).

Active egress of *Toxoplasma* from host cells requires rupture of the parasitophorous vacuole membrane (PVM) and the host cell membrane (9). Egress is essential for the dissemination of the infection and it has been known for several years that Ca^2+^ ionophores can trigger egress (10). However, it was the use of GECIs that provided the final and conclusive evidence that there was a cytosolic Ca^2+^ increase right before egress (7). Secretion of the perforin-like protein 1 (TgPLP1) from micronemes (specialized secretory organelles involved in egress, motility, and invasion by tachyzoites), assists in the permeabilization of the PVM and host cell membrane (11). Both secretion of the microneme protein TgPLP1 and initiation of motility during egress are stimulated by an increase in cytosolic Ca^2+^. It has been proposed that the trigger for this cytosolic Ca^2+^ increase is the rupture of the host plasma membrane and the ensuing reduction in the concentration of cytoplasmic potassium. Low [K^+^] would activate a phospholipase C activity in *Toxoplasma* that, in turn, would cause an increase in cytoplasmic [Ca^2+^] in the parasite (12).

As an obligate intracellular parasite, *T. gondii* resides and replicates within the PV that functions as a molecular sieve to passively permit the exchange of small molecules; thus, the surrounding milieu of intracellular parasites is likely in equilibrium with the host cell cytoplasm (13). Therefore, intracellular parasites are likely exposed to the fluctuations of the host cytosolic ionic composition. The host cytoplasmic Ca^2+^ level is highly controlled and kept at physiological levels of ∼70-100 nM under basal conditions, which is similar to the cytoplasmic levels of the replicating parasites. Experimental evidence indicates that Ca^2+^ efflux from intracellular stores represent the first initial step of the Ca^2+^ signaling pathway for egress, so maintaining the high Ca^2+^ levels of the stores is fundamental for continuation of the lytic cycle.

In this work, we investigated both the ability of *Toxoplasma* to replenish its intracellular Ca^2+^ stores during its replication and the role of low [K^+^] as a trigger for host cell egress. Using a variety of pharmacological tools, fluorescence microscopy, and a novel approach using patched host cells, we show that Ca^2+^ signaling of the host cell impacts parasite Ca^2+^ levels and drives parasite egress, while low [K^+^] modulates but is not the trigger for parasite egress.

## Results

### Calcium influx in intracellular parasites

We previously characterized a Ca^2+^ influx pathway at the plasma membrane of extracellular *T. gondii*. Following on this finding we wanted to determine if Ca^2+^ influx was also operational in intracellular replicating parasites. For this we measured cytosolic Ca^2+^ responses of intracellular tachyzoites and exposed them to fluctuations of host cytosolic Ca^2+^ by stimulating them with a variety of agonists that act specifically on the host cell. We used Genetically Encoded Ca^2+^ Indicators (GECIs) (14, 15) expressed in the cytosol of HeLa cells infected with *T. gondii* tachyzoites expressing either cytosolic GCaMP6f or luminal PV targeted jGCaMP7f (16) (See Material and Methods and Table S1). We grew HeLa cells on coverslips, transfected them with red GECIs and infected these cells with green GECI-expressing tachyzoites. To follow Ca^2+^ changes, we used time-lapse microscopy and stimulated them with specific host cell agonists (Figs 1 and S1). We first used carbachol, a muscarinic receptor agonist that specifically acts on mammalian host cells and stimulates Ca^2+^ oscillations (17). Muscarinic receptors, stimulated by the neurotransmitter acetylcholine, are well-characterized G-coupled protein receptors that lead to an increase in cytosolic Ca^2+^ via activation of a phosphatidyl inositol phospholipase C (PI-PLC) and generation of inositol 1,4,5-trisphosphate (IP_3_) (18). Carbachol addition resulted in a dramatic rise in the fluorescence of RGECO or jRGECO1a, indicating an increase in host cytosolic Ca^2+^ (Fig 1A, *red tracings*). For the experiment shown in Fig 1A, the tachyzoites expressed jGCaMP7f in the PV, which was achieved by fusing the *T. gondii* P30 gene to the N-terminus of the GECI, which was shown to confer localization to the PV (11). An almost simultaneous rise in PV Ca^2+^ and host Ca^2+^ was observed (Fig 1A, *green tracing*) thereby supporting the molecular sieve model of the PV (13). We also tested if the parasite cytosolic Ca^2+^ also increased following host cytosolic Ca^2+^ stimulation thus supporting Ca^2+^ influx from the host cytosol to the parasite. HeLa cells expressed a cytosolic copy of RGECO and were infected with tachyzoites expressing GCaMP6f in their cytoplasm (Fig 1B, *green tracing* and Supplemental Video 1). Ca^2+^ influx into the intracellular tachyzoite was also observed in experiments using HeLa cells infected with parasites expressing GCaMP6f in their cytosol and RGECO localized to the lumen of the PV. Carbachol addition caused a uniform rise in all PVs expressing the GECI, followed by a small proportion of parasites showing increase in cytosolic Ca^2+^ (GCaMP6f signal) (Fig 1C). It is possible that this phenotype represents parasites at different stages of the replication cycle that are more or less sensitive to Ca^2+^ influx. All parasites within the same PV were exposed to the same rise in luminal PV Ca^2+^ because all PV’s responded to carbachol stimulation (Fig 1A), yet only a few parasites showed a subsequent increase in parasite cytosolic Ca^2+^ (Fig 1B).

**Figure 1:**
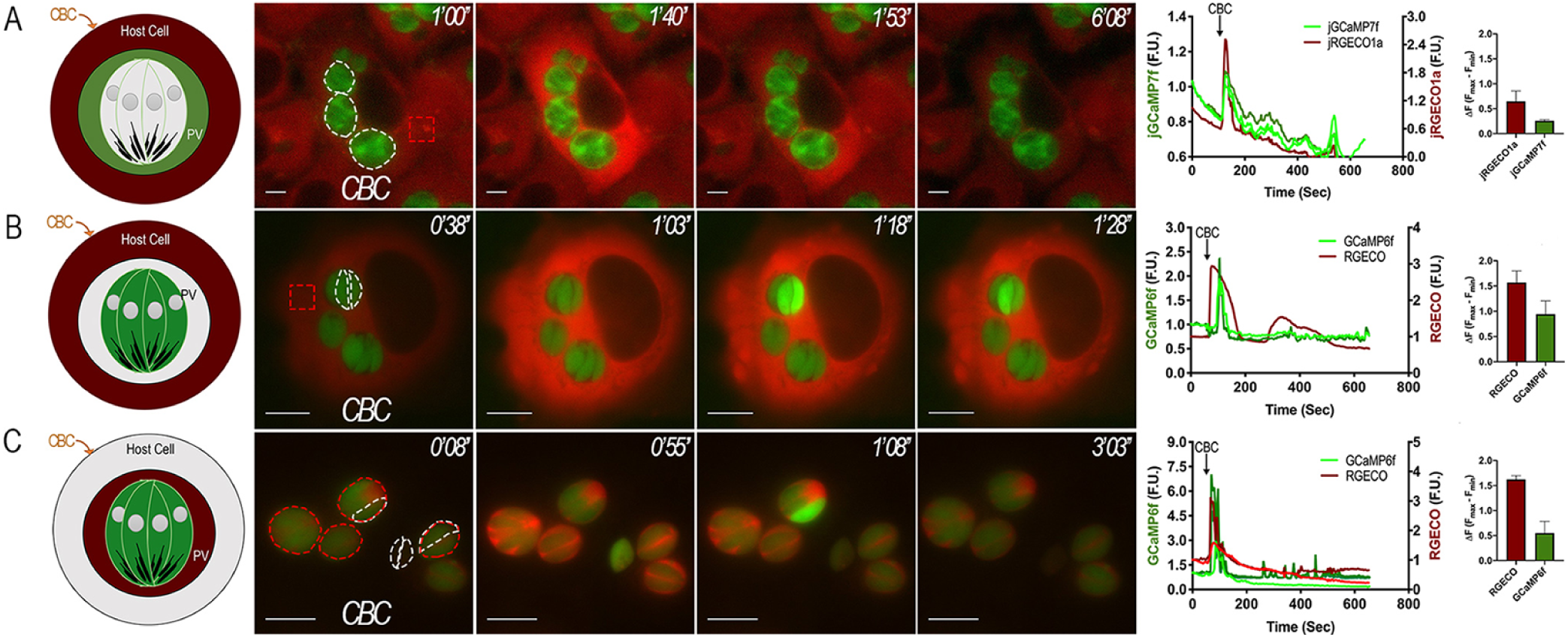
Ca^2+^ Host Signaling and *T. gondii* egress: A) Representative Images of Hela cells transiently expressing the red GECI RGECO that were infected with *cyto*-GCaMP6f expressing parasites. 1 mM Carbachol was added 1 min after recording started. B) Representative Images of Hela cells stably expressing jRGECO1a that were infected with PV localized P30-jGCaMP7f expressing tachyzoites. 1 mM Carbachol was added 1 min after recording started. C) Representative images of Hela cells infected with P30-RGECO *cyto*-GCaMP6f. 1 mM Carbachol was added 1 min after recording started Tracings to the right of each panel shows fluorescence fluctuations of single parasites (*green*) or of a delineated region of interest in the host cells (*red*) or PV (green) from 3 independent trials for each condition. Bar graphs represent quantification of the average ΔF values of three independent trials (Red, jRGECO1a or RGECO, *right axis*) (*Green*, GCaMP6f or jGCaMP7f, *left axis*). Dashed white outlines indicate the area used as a region of interest for the analyzes for fluorescence changes. Dashed white outlines show GCaMP6f-expressing parasites whose fluorescence tracings were used for analysis. Dashed red outlines indicate the region of the host cell that was used to analyze the RGECO or jRGECO1a channel. Numbers at the upper right of each panel indicate the time frame of the video.

We repeated our experiments assessing the Ca^2+^ waves between host-PV-parasite cytosol by stimulating host Ca^2+^ increase using histamine (Fig S1). Histamine functions through a similar, though different G-coupled protein receptor than carbachol. Histamine will also induce an IP_3_ mediated increase in cytosolic Ca^2+^ (19). The results with histamine were similar to the ones with carbachol in which a rise in host cell Ca^2+^ occurred simultaneous with PV Ca^2+^. Additionally, parasite Ca^2+^ influx trailed host Ca^2+^. These results demonstrate that intracellular tachyzoites are able to take up Ca^2+^ from the host Ca^2+^ cytosol following Ca^2+^ signaling events.

### A Threshold of Ca^2+^ is needed for Parasite Egress

To visualize the link between host Ca^2+^ signaling and intracellular parasite Ca^2+^ fluctuations and egress, we transiently transfected HeLa cells with LAR-GECO1.2 (a red Ca^2+^ indicator optimized for expression within the host mitochondria) (20) and infected the host cells with cyto-GCaMP6f expressing tachyzoites. We stimulated Ca^2+^ with ionomycin, a Ca^2+^/H^+^ ionophore which causes Ca^2+^ release from all neutral stores (21), or thapsigargin (TG), an inhibitor of the endoplasmic reticulum (ER) Ca^2+^-ATPase SERCA (Fig 2A&B) (22). The ER constantly allows efflux of Ca^2+^ via an unknown mechanism, and the SERCA serves as a counterbalance to pump Ca^2+^ back into the organelle. Blocking of SERCA via thapsigargin will induce a rise in cytosolic Ca^2+^ due to the ER’s constant efflux of Ca^2+^ (22). The addition of ionomycin leads to a dramatic Ca^2+^ increase in both host cell and parasite, followed by rapid egress (Fig 2A&C). TG induced an increase in the fluorescence of LAR-GECO, indicating an increase of Ca^2+^ within the host mitochondria (LAR-GECO), as well as Ca^2+^ oscillations within the parasite (GCaMP6f). The increase in host mitochondrial Ca^2+^ can be accounted for as due to the close, tight interaction between the ER and the mitochondria (23). Interestingly, addition of TG caused Ca^2+^ oscillations in the intracellular tachyzoites but they remained intracellular throughout all 10 min of recording (Fig 2B&D and Supplemental Video 2).

**Figure 2:**
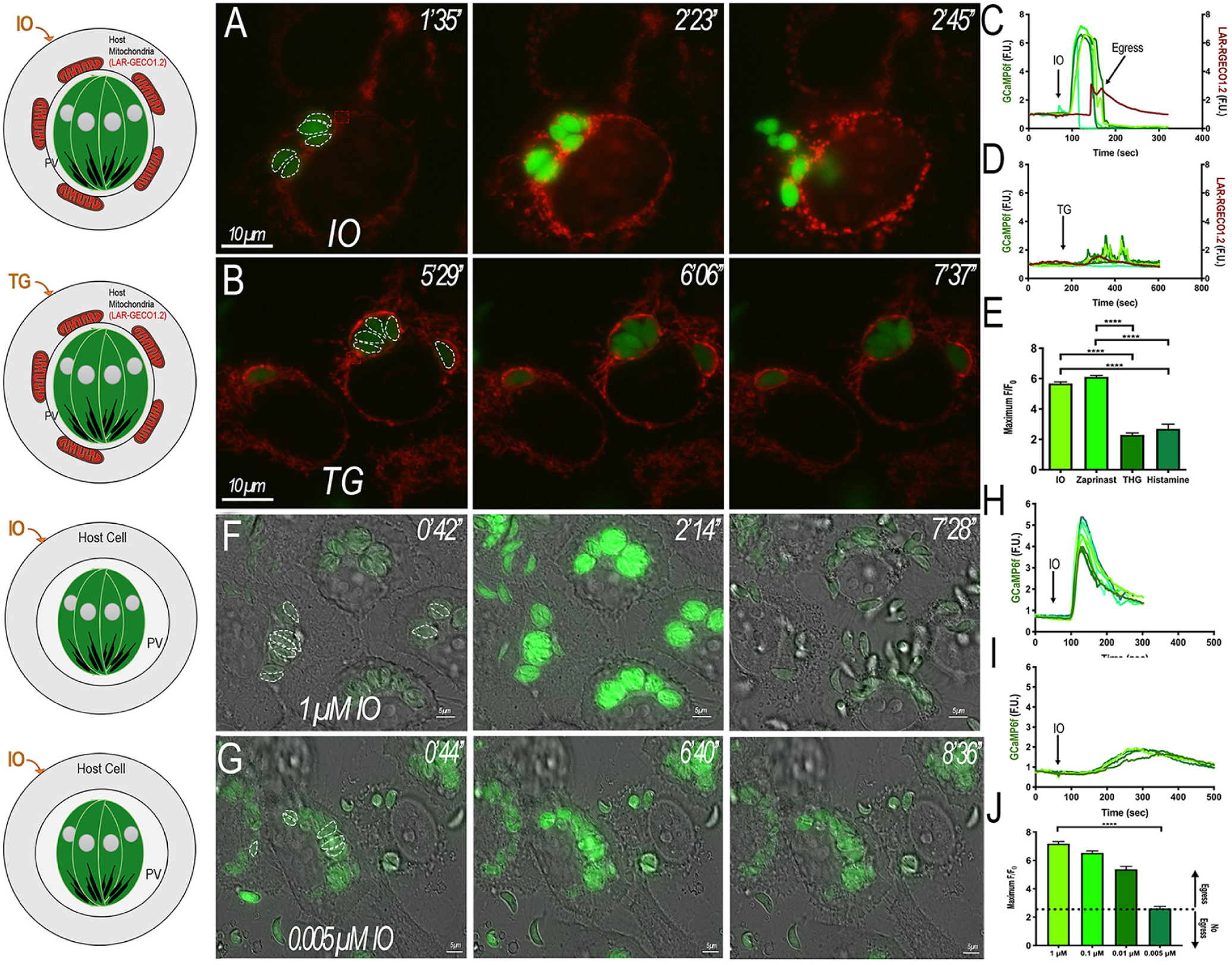
Threshold for Egress: A & B) Hela cells were transiently transfected with the low affinity mitochondria Ca^2+^ indicator LAR-GECO1.2 and infected with tachyzoites expressing cyto-GCaMP6f for approximately 20 h. The mitochondria surrounding each PV is labeled with the LAR-GECO1.2 (*red*). 1 µM Ionomycin (IO) and 2 µM Thapsigargin (TG) were added, respectively at 1 min and the response recorded. Dashed white outlines indicate the region of interest used for the analysis of the fluorescence changes. Dashed white outlines show GCaMP6f-expressing parasites whose fluorescence tracings were used for analysis. Dashed red outlines indicate the region of the host cell that was used to analyze for the LAR-GECO1.2 channel. Numbers at the upper right of each panel indicate the time frame of the video. C & D) Fluorescence tracings obtained after addition of IO or TG, respectively. Dashed white outlines shown in A&B indicate parasites used for analysis of GCaMP6f parasites. Dashed red outlines indicate region of LAR-GECO1.2 channel that was used for analysis. Numbers at the upper right of each panel indicate the time frame of the video. E) Comparison of Δ-Fluorescence values from each Pharmacological stimulus. IO and Zaprinast (Zap) induce egress while TG and Histamine did not. F & G) Hela cells were infected with GCaMP6f parasites and IO was titrated down to find a concentration that did not induce egress (0.005 µM). Dashed white outlines indicate the area used as a region of interest to analyze the fluorescence changes. Numbers at the upper right of each panel indicate the time frame of the video. H & I) Fluorescence tracings of IO threshold titration of 1 µM and 0.005 µM, respectively. J) Average Δ-fluorescence values of 1, 0.1, 0.01, and 0.005 µM IO.

We next tested Zaprinast, a cGMP phosphodiesterase inhibitor, which inhibits the cGMP phosphodiesterase resulting in a build-up of cGMP that activates protein kinase G (PKG) leading to an increase of cytosolic Ca^2+^ and stimulation of egress (24, 25). The increase of cytosolic Ca^2+^ by Zaprinast was almost comparable to the increase observed with ionomycin, and also resulted in egress (Fig 2E). However, while both histamine and thapsigargin did induce an increase in parasite cytosolic Ca^2+^, the response was insufficient to induce egress. Ionomycin and Zaprinast produced a ΔF_max_/F_0_ of approximately 6 that was not statistically different between the two. Thapsigargin or histamine ΔF_max_/F_0_ responses were much lower of approximately ∼2 fold and was not statistically different between the two reagents. Comparing the results with Ionomycin to those with either thapsigargin or histamine and the results with Zaprinast to those of either thapsigargin or histamine, show that the responses were statistically different (Fig 2E). Our results suggest that a threshold for cytosolic Ca^2+^ has to be met in order to induce egress.

With the aim of estimating the Ca^2+^ threshold needed, we titrated down ionomycin to determine a concentration that would still produce a Ca^2+^ response but was insufficient to induce egress. Ionomycin at 1 μM induced a rapid egress response (Fig 2F&H), and titrating down the concentration of ionomycin to 0.1 µM and 0.01 µM led to a decrease in the ΔFluorescence response, though parasite egress was still observed. No egress was observed when the ionomycin concentration was lowered to 0.005 µM (Fig 2G&I). It is significant to note that the ΔFluorescence response of 0.005 μM was approximately 3, a value closely similar to the responses of thapsigargin and histamine that also did not produce a sufficient response to induce egress (Fig 2E&J). The cytosolic Ca^2+^ concentration that tachyzoites reached in response to various concentrations of Ionomycin were evaluated in a separate experiment with extracellular tachyzoites loaded with the ratiometric Ca^2+^ indicator Fura-2-AM (Fig S3). The cytosolic concentration reached for Ionomycin at 5 nM was ∼250 nM and at 10 nM was around 750 nM. From these experiments we could conclude that there is a cytosolic Ca^2+^ threshold that needs to be reached for the stimulation of egress and it is around 300-500 nM.

### Two Peaks of Ca^2+^ Lead to Parasite Egress

To further characterize the Ca^2+^ response of intracellular parasites and the ensuing egress we tested Zaprinast, which will result in Ca^2+^ increase and stimulation of microneme secretion, and parasite egress. This experiment was done in the presence of extracellular buffer supplemented with 2 mM Ca^2+^ and we noticed that egressing parasites displayed a quantifiable, characteristic two fluorescence peaks that preceded egress (Fig 3A-C and Supplemental Video 3). We repeated the Zaprinast induced egress experiments with extracellular buffer supplemented with 100 µM EGTA (Ca^2+^ free) and quantified the response. Under these conditions the second fluorescence peak was lower and wider (Fig 3D-F). Parasites took longer to egress when Zaprinast was added in Ca^2+^ free buffer compared to egress in extracellular buffer supplemented with 2 mM Ca^2+^ (Fig 3F).

**Figure 3:**
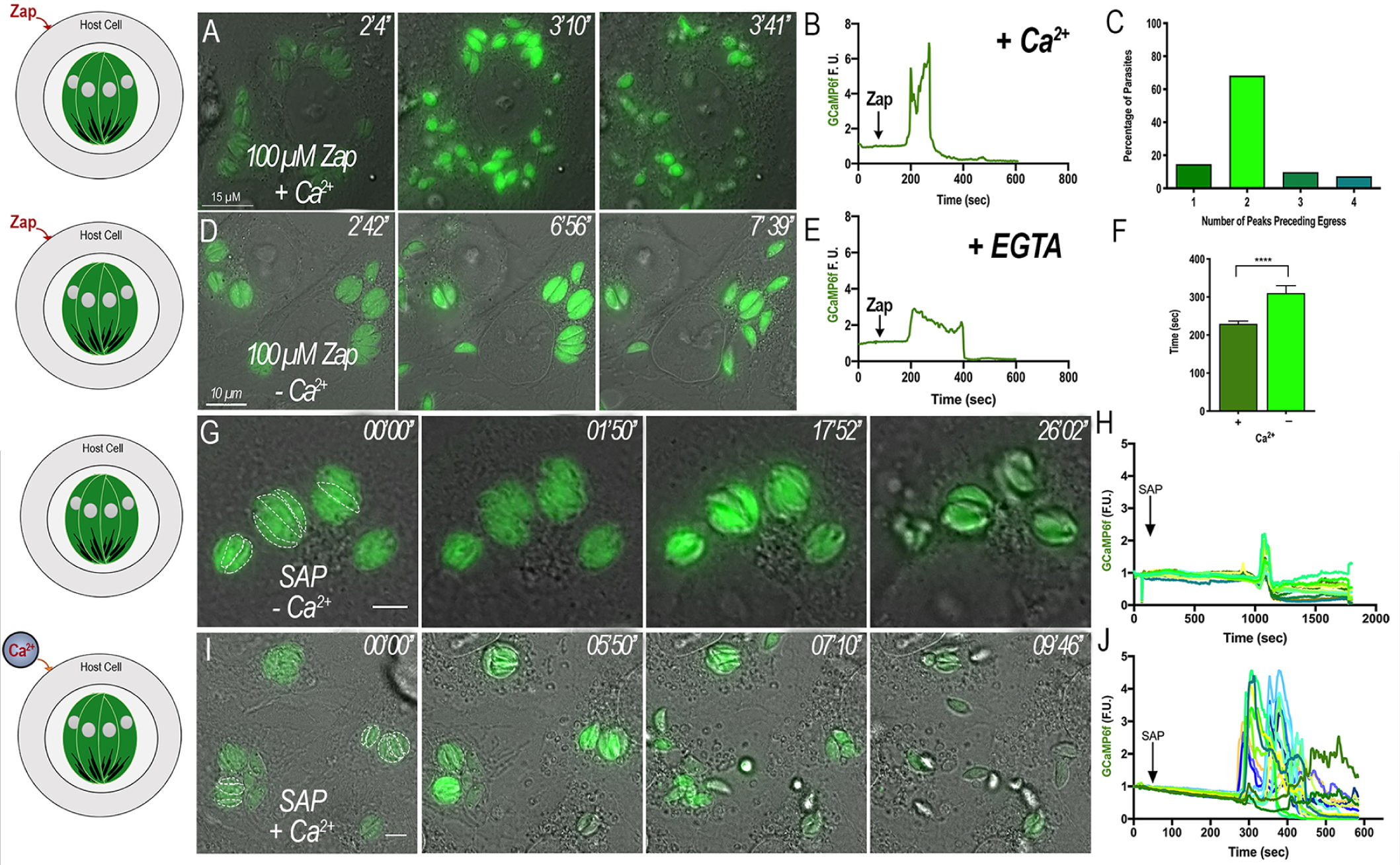
Two Peaks of Ca^2+^ Precede Egress: A & D) Hela cells infected with GCaMP6f parasites bathed in Ringer Buffer supplemented with either 2 mM Ca^2+^ (Ca^2+^ rich) or 100 µM EGTA (Ca^2+^ free media), respectively post addition of Zaprinast. Dashed white outlines show the region of interest for the analysis of the fluorescence changes shown in the tracings. Numbers at the upper right of each panel indicate the time frame of the video. B & E) Representative fluorescence tracings of single parasites under Zaprinast simulation in the presence and absence of extracellular Ca^2+^, respectively. Under Ca^2+^-free conditions (100 μM EGTA) the second peak is reduced. C) Quantification of GCaMP6f peak number preceding egress. Graph represents the summation of 3 independent experiments. F) Average time of egress comparing Zaprinast induced egress resuspended in extracellular media supplemented with either 2 mM Ca^2+^ (+) or 100 μM EGTA (-). G) Intracellular GCaMP6f parasites were exposed to the surrounding buffer under Ca^2+^ free conditions (1 mM EGTA) with a 0.01% (w/v) saponin solution in Ringer Buffer (SAP – Ca^2+^). Dashed white outlines represent the parasites that were used in analysis. Numbers at the upper right of each panel indicate the time frame of the video. H) Representative fluorescence tracings of non-egressing parasites treated with saponin in Ca^2+^-free conditions as shown in G. I) Intracellular GCaMP6f parasites were exposed to the surrounding buffer under Ca^2+^ rich conditions (2 mM Ca^2+^) using a 0.01% (w/v) saponin solution in Ringer Buffer (SAP + Ca^2+^). Dashed white outlines represent the parasites that were used for the analysis. Numbers at the upper right of each panel indicate the time frame of the video. J) Representative fluorescence tracings of egressing parasites treated with saponin in Ca^2+^-rich conditions as shown in J.

We next compared the parasite cytosolic Ca^2+^ fluctuations after permeabilizing the host cell with saponin under two conditions: Ca^2+^ free (100 μM EGTA) and Ca^2+^ rich (2 mM Ca^2+^) buffers (Fig. G-J). Without extracellular Ca^2+^, egress was not observed, though a sharp single peak was evident after quantifying the fluorescence tracings (Fig 3G-H). In comparison, when using a Ca^2+^ rich buffer after permeabilizing the host membrane to expose the parasites to the extracellular buffer, cytosolic Ca^2+^ levels oscillated and they promptly egressed from their respective PV’s (Fig 3I-J). In the presence of extracellular Ca^2+^ two peaks of Ca^2+^ were observed (Fig. 3J).

The microneme protein Perforin-Like Protein 1 (PLP1) functions in breaking down the host cell during egress, exposing intracellular tachyzoites to the surrounding extracellular milieu. PLP1 has been shown to be involved in the initial rupture of the PV (11) and the release of Ca^2+^ from intracellular stores could stimulate this release in the natural egress process. Our model proposes that this rupture would allow the extracellular Ca^2+^ to influx the parasites contributing to reaching the threshold for downstream stimulation of gliding motility and egress. We tested whether ΔPLP1 mutants, defective in their ability to breakdown the host cell, would still display two peaks during egress. We transfected GCaMP6f into ΔPLP1 mutants and induced egress using 100 μM Zaprinast (Fig S2). We quantified the fluorescence tracings of egressing parasites and these cells only displayed 1 peak (Fig S2B &C) supporting that the second peak is due to influx from the extracellular milieu.

These experiments demonstrate that intracellular tachyzoites are capable of taking up Ca^2+^ from the host cytosol and also from the extracellular milieu after host cell rupture which results in a second peak of cytosolic Ca^2+^ that precedes egress.

### A Ca^2+^ Peak Precedes Natural Egress from Host Cells

With the aim of investigating the presence of two peaks of cytosolic Ca^2+^ during natural egress we synchronized intracellular parasites using compound 1 (cpd1) (Fig 4 and Supplemental Video 4). In the related Apicomplexa parasite *Plasmodium falciparum*, cpd1 reversibly inhibits PKG and arrests parasites from egressing (26). We pre-incubated HeLa cells expressing jRGECO1a and infected with GCaMP6f-expressing tachyzoites with cpd1 for 24 h to arrest egress. At this time, we washed off cpd1 and egress was observed within approximately two min preceded by an increase in cytosolic Ca^2+^. The increase began with a single “leader” tachyzoite that was distinguishable due to having the largest rise in cytosolic Ca^2+^ and this cell egressed first (Fig 4 A). We quantified the fluorescence tracings, and as expected, egressing parasites displayed two peaks during egress (Fig 4 C-E). Interestingly, a rise in host Ca^2+^ was also evident during the natural egress process, thus highlighting the role of extracellular Ca^2+^ influx during natural egress (Fig. 4C&D, *red tracing*). Next we tested how ΔPLP1 parasites would respond under natural egress conditions (Fig. 4B&F). Post cpd1 washout, ΔPLP1 GCaMP6f oscillated randomly and nonuniformly, and no parasite exhibited the two-peak pattern observed in wild type parasites (Fig. 4F). Some parasites displayed movement within the PV of the ΔPLP1 parasites(11) post rise in cytosolic Ca^2+^ though none of these parasites displayed a second peak of higher amplitude that would be associated with Ca^2+^ influx, as well as no rise in the jRGECO1a channel that would be indicative of rupture of the host cell and extracellular Ca^2+^ influx. Eventually the fluorescence of these parasites diminished indicating lower cytosolic Ca^2+^, and the parasites stopped moving shortly after. Given these results, we conclude that two-peaks of Ca^2+^ are part of the natural egress progression of *T. gondii* and the second peak is due to Ca^2+^ influx from the extracellular milieu.

**Figure 4:**
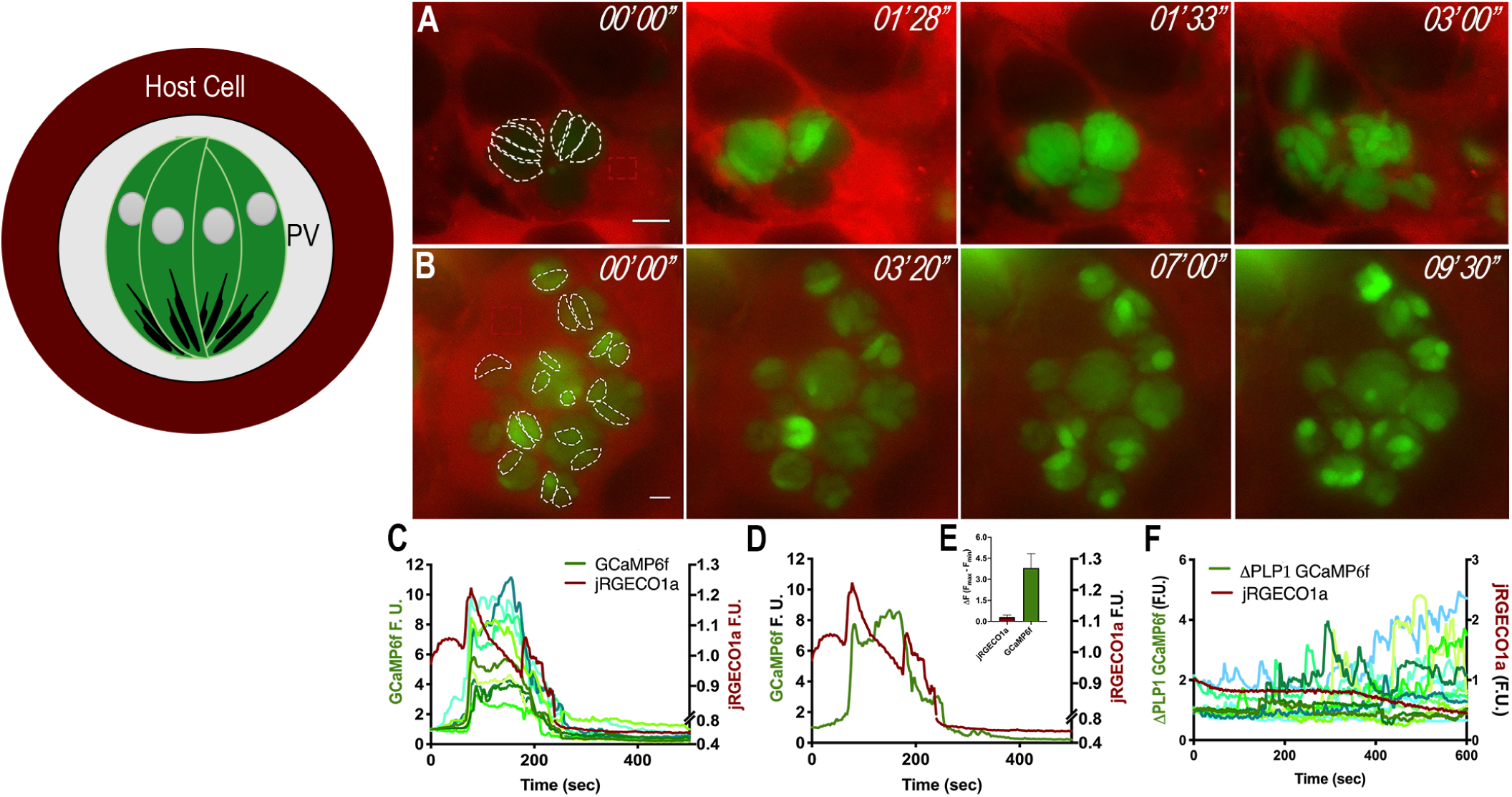
Natural Egress: Intracellular GCaMP6f parasites were synchronized for natural egress with Compound 1. A) Still images of Hella cells expressing jRGECO1a infected with tachyzoites expressing GCaMP6f and treated with 1 μM compound for 24 hours. After washout of compound 1, parasites egressed naturally within 3-5 mins; dashed regions indicate the area used as region of interest for the analysis of fluorescence changes shown in the tracings. Dashed white outlines are used to highlight GCaMP6f-expressing parasites whose fluorescence tracings were used for analysis. Dashed red outlines indicate the region of the host cell that was used to analyze for the jRGECO1a channel. Numbers at the upper right of each panel indicate the time frame of the video. B) Still images of Hella cells expressing jRGECO1a infected with ΔPLP1 tachyzoites expressing GCaMP6f and treated with 1 μM compound for 24 hours. After washout of compound 1, parasites egressed naturally within 3-5 mins; dashed regions indicate the area used as region of interest for the analysis of fluorescence changes shown in the tracings. Dashed regions are used to highlight ΔPLP1 GCaMP6f-expressing parasites whose fluorescence tracings were used for analysis. Dashed red outlines indicate the region of the host cell that was used to analyze for the jRGECO1a channel. Numbers at the upper right of each panel indicate the time frame of the video. C) Fluorescence tracings of host cell jREGO1a (red tracing) and GCaMP6f parasites (green tracings); D) Single Fluorescence tracings of host cell jREGO1a (red tracing) and GCaMP6f parasites (green tracings). Note that after compound 1 washout parasites egress still occurs with two peaks of fluorescence; E) Quantification of ΔF of jRGECO1a (red bar) and GCaMP6f (green bar) after cpd1 washout. F) Fluorescence tracings of host cell jREGO1a (red tracing) and ΔPLP1 GCaMP6f parasites (green tracings)

### Blocking Ca^2+^ Influx Using Pharmacological Agents Blocks Parasite Egress

With the aim of characterizing further the source of the two peaks of Ca^2+^ we tested some pharmacological agents. We first tested nifedipine, a voltage-gated Ca^2+^ channel inhibitor and pretreated intracellular cyto-GCaMP6f-expressing parasites with the drug to compare egress with control parasites (Fig. 5A). The surrounding extracellular buffer was supplemented with 2 mM Ca^2+^ and the host cells were permeabilized with saponin. While the control non-treated parasites egressed following a sharp rise in cytosolic Ca^2+^ trailed by smaller Ca^2+^ oscillations (Fig 5A &B) the nifedipine treated parasites did not egress, and showed only modest Ca^2+^ oscillations of approximately two-fold range (Fig 5C&D), which are below the threshold needed for egress (Fig 2). We next tested cpd1 to inhibit PKG and induced egress with Zaprinast. While control parasites egressed following a fast rise in cytosolic Ca^2+^ with two peaks (Fig 5E&F), pretreatment with cpd1 abolished egress, and the parasites’ cytosolic Ca^2+^ only raised about 3-fold (Fig 5G&H). These results showed that blocking extracellular Ca^2+^ influx with nifedipine or inhibiting PKG, results in intracellular parasites that are unable to reach the threshold needed for egress, in support for the role of extracellular Ca^2+^ influx in egress.

**Figure 5:**
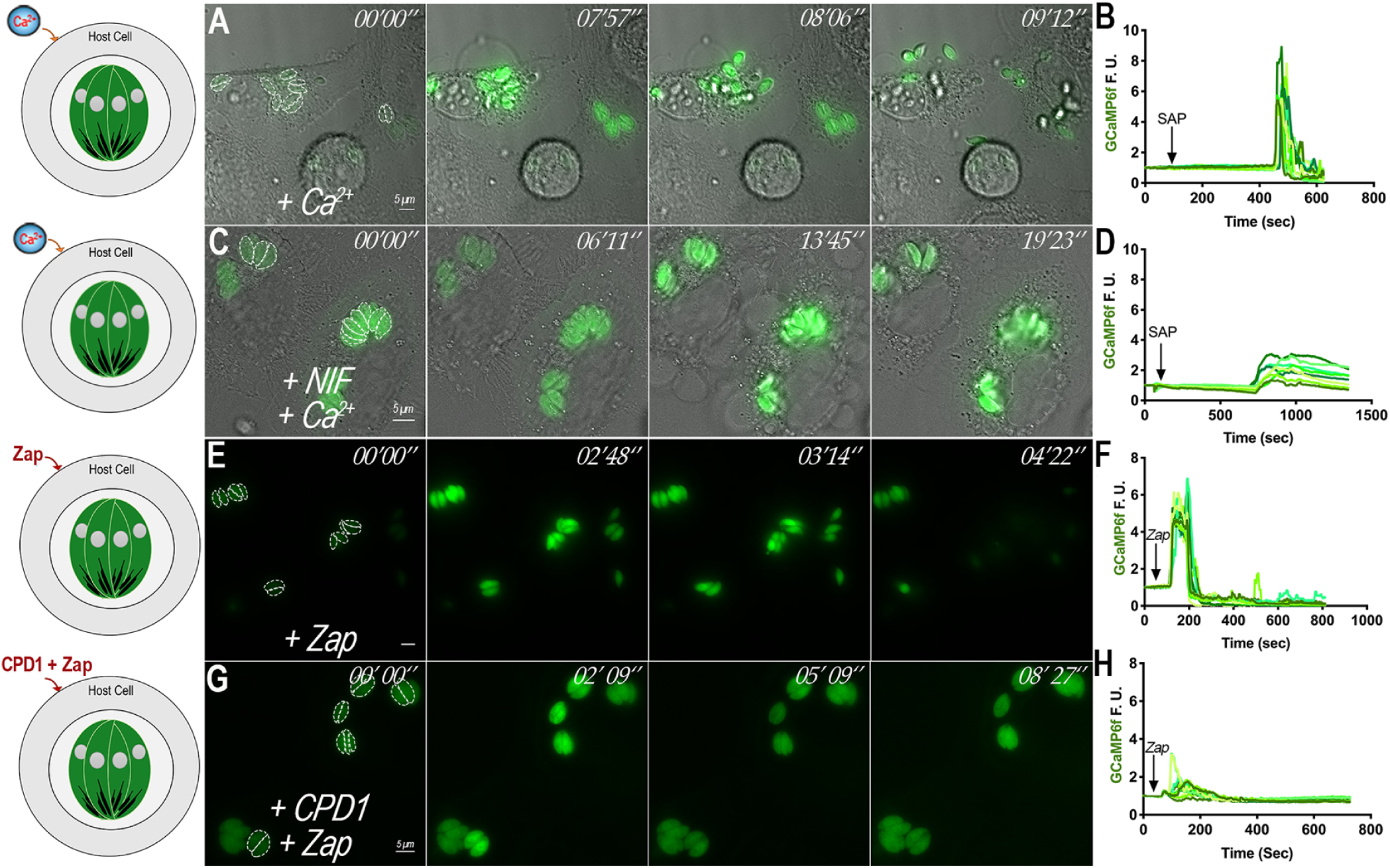
Pharmacological inhibition of Ca^2+^ influx leads to Inhibition of Egress: A) GCaMP6f infected host cells were exposed to a buffer supplemented with 2 mM Ca^2+^ containing 0.01% (w/v) saponin, and fluorescence changes were recorded. B) Fluorescence tracings of egressing parasites after addition of saponin in media supplemented with 2 mM Ca^2+^; C) Hela cells infected with tachyzoites expressing GCaMP6f were pretreated for 5 min with 10 μM Nifedipine, a L-type Voltage Gated Calcium Channel Blocker in the presence of an extracellular buffer supplemented with 2 mM Ca^2+^ containing 0.01% (w/v) saponin and fluorescence changes were recorded. Note that parasites did not egress under these conditions; D) Fluorescence tracings of GCaMP6f parasites pretreated with Nifedipine after saponin permeabilization. E) Hela cells infected with GCaMP6f expressing tachyzoites were treated with 100 μM Zaprinast and egressed rapidly afterwards; F) Fluorescence tracings of GCaMP6f parasites treated with 100 μM Zaprinast as indicated by the dashed outlines in E. G) Hela cells infected with GCaMP6f expressing tachyzoites were pretreated with 1 μM of compound 1 for 5 mins. After addition of 100 μM Zaprinast, parasite cytosolic Ca^2+^ slightly and briefly oscillated but no egress was evident; H) Fluorescence tracings after pretreatment with 1 μM compound 1 and stimulation of egress with 100 μM Zaprinast as indicated by the dashed white outlines in G. dashed outlines indicate the area used as a region of interest for the analysis of the fluorescence changes shown in the tracings. For A, C, E and G, the white outlines highlight the GCaMP6f-expressing parasites whose fluorescence tracings were used for analysis presented in B, D, F and H. Numbers at the upper right of each panel indicate the time frame of the video.

### Role of K^+^ and Ca^2+^ in parasite egress

The cytosol of eukaryotic cells contains approximately 140 mM K^+^ with 70-90 nM Ca^2+^, while the extracellular milieu contains 5 mM K^+^ and ∼1.8 mM Ca^2+^. During a Ca^2+^ signaling event, cytosolic Ca^2+^ rises rapidly to μM levels (5), and we have shown that a crosstalk exists between the host cell and the parasite cytosolic Ca^2+^ (Fig 1). To manipulate the host cytosolic ionic composition, and identify the Ca^2+^ concentration that would induce parasite egress, we patched infected HeLa cells (27). We attached a pipet to the plasma membrane of infected cells as shown in Fig 6A, formed a patch seal between the plasma membrane and the pipette, and broke the plasma membrane, thus equilibrating the solution in the patch pipette with the cytosol of the host cell. This technique allowed us to control the composition of the buffer surrounding the PVs by modifying the pipet buffer (Fig 6A). We tested increasing concentrations of cytosolic free Ca^2+^ (0.5, 1, 5, and 10 μM) in a 140 mM K^+^ patch pipette solution (Fig. 6B). We did not observe egress at 0.5 μM Ca^2+^, and only a small percentage of parasites egressed at 1 μM Ca^2+^. Approximately one third of parasites egressed when exposed to 5 μM Ca^2+^ while the remaining two thirds did not egress. Ultimately at a concentration of 10 μM Ca^2+^, all parasites egressed (Supplemental Video 5). Previous literature has stated that the high potassium concentration of the host cytosol prevented parasite egress. It was postulated that a decrease in the concentration of K^+^ would activate a Ca^2+^ signal, inducing microneme secretion and subsequent egress (12). To test the role of K^+^ in parasite egress, we repeated the whole-cell patch in a low potassium solution within the patch pipette (Fig 6C). Choline chloride was added to maintain the same osmolarity and anionic composition. Under low potassium, we did not observe egress at 0.5 μM Ca^2+^, and two thirds of the parasites of the patched cells egressed at 1 μM Ca^2+^, and parasites egressed faster compared to their high potassium counterpart. Approximately 80% of parasites egressed at 5 μM Ca^2+^, and egress occurred in approximately 3 min compared to 5 min in high K^+^. Similar to the high potassium conditions, 10 μM Ca^2+^ caused 100% egress, but again at a much faster rate. We summarized our patch data in Fig. 6D&E. No egress was observed under both conditions for 0.5 μM Ca^2+^, for both conditions and percentage and rate of egressing parasites was higher and faster under low potassium conditions. As we increased the cytosolic Ca^2+^ concentration to 2, 5, and 10 μM Ca^2+^, the percentage of egressing parasites increased quasi-linearly under the low potassium conditions. Under high potassium conditions the percentage of egressing parasites increased quasi-linearly from 1-5 μM Ca^2+^, but 10 μM Ca^2+^ was sufficient to induce 100% egress (Supplemental Video 6). At 1 μM Ca^2+^, parasites egressed in approximately 550 and 320 sec for high and low potassium, respectively. In high potassium conditions, parasites egressed at about the same rate for 2 and 5 μM Ca^2+^, but under low potassium conditions parasites egressed faster at 5 μM Ca^2+^ versus 2 μM Ca^2+^ (∼300 seconds (2 μM Ca^2+^) versus ∼200 seconds (5 μM Ca^2+^)). Though 100% egress was observed at 10 μM Ca^2+^ for both conditions, in high potassium parasites egressed in ∼250 seconds, versus ∼50 seconds in low potassium conditions. In summary, according to these results Ca^2+^ is crucial for egress but a decrease in the concentration of K^+^ as the parasite lyses the host cell and egresses, results in acceleration but is not the trigger for egress.

**Figure 6:**
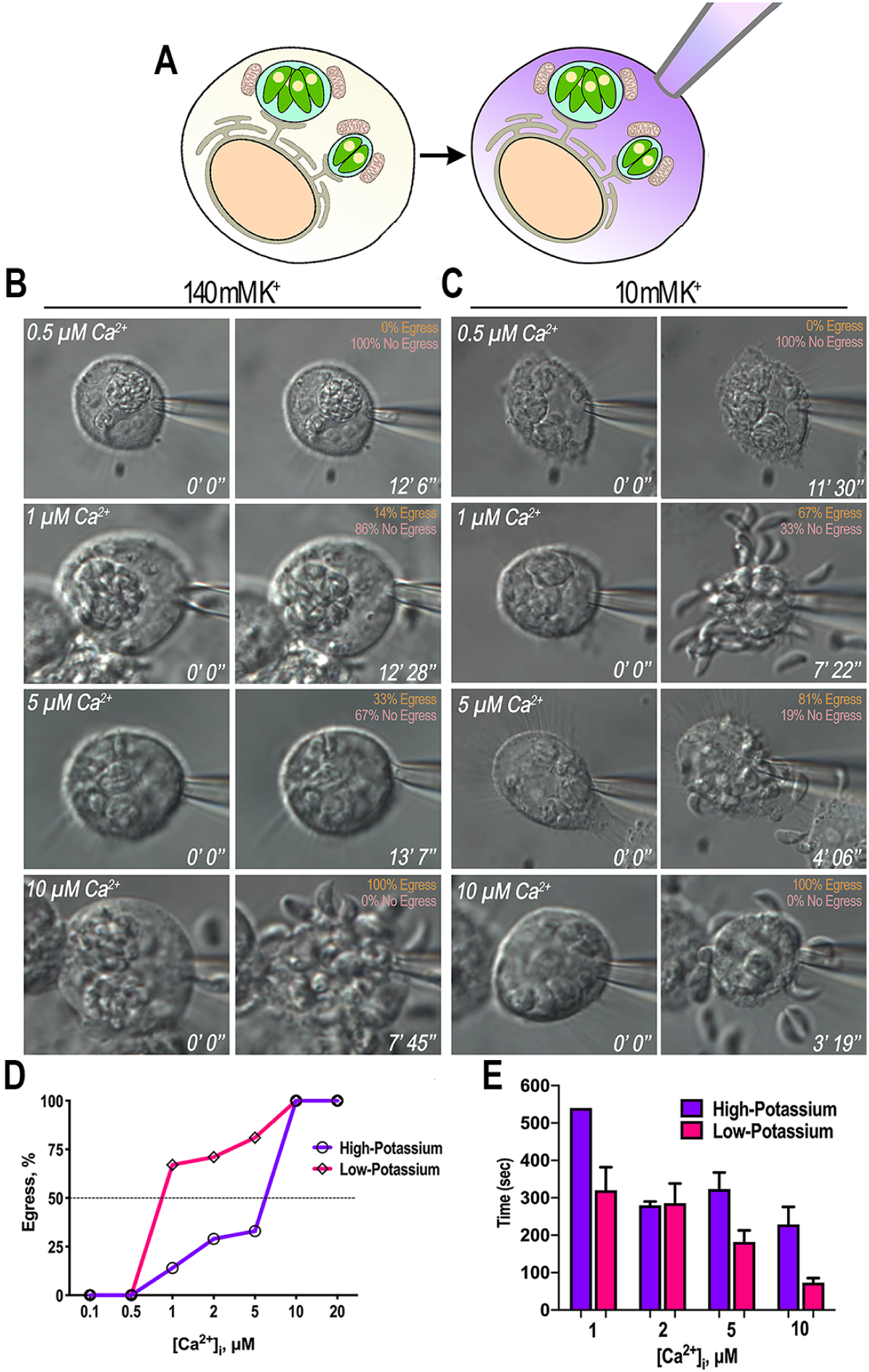
Host-Cell Patch: Hela cells infected with GCaMP6f tachyzoites were whole-cell patched and egress was monitored. A) Whole-cell patch allowed for the exposure of PVs to defined Ca^2+^ concentrations by exchanging the cytosol of the host cell with the composition of the buffer inside the patch pipette; B) Representative still images of infected host cells were patched under high potassium conditions (140 mM K^+^) to mimic the high potassium environment of the host cytosol. Gradually increasing concentrations of free calcium were tested to see if/when egress occurred. The percentage of egressing vs non-egressing parasites is shown in the upper left-hand corner; C) Representative still images of infected hosts cells that were patched under low potassium conditions (10 mM K^+^ and 130 mM Choline Chloride) and egress was monitored under the same experimental conditions as in A; D) Percentage of egressing parasites as a function of Ca^2+^ concentration. Green, 140 mM K^+^, Red, 10 mM K^+^ and 130 mM K^+^; E) Average time of egress under high (140 mM K^+^) and low potassium conditions (10 mM K^+^ and 130 mM Choline Chloride). Note that under low K^+^ conditions the percentage of egressing parasites increases, and parasites egress faster.

## Discussion

Our work shows that Ca^2+^ influx into the parasite during host physiological Ca^2+^ signaling events (i.e. stimulation of GPCR, VGCC, etc) could be the means the parasite uses to replenish its intracellular Ca^2+^ stores depleted after host cell invasion. Intracellular tachyzoites via the PV are exposed to the low Ca^2+^ concentration (70-100 nM) present in the host cytosol while extracellular tachyzoites are surrounded by high Ca^2+^ (1.8-2.0 mM). Considering that egress can occur at nearly any stage of intracellular replication and is preceded by a spike in cytosolic Ca^2+^ (7), that very likely is initiated by intracellular store release, intracellular replicating *T. gondii* needs to maintain its intracellular stores filled with Ca^2+^ (7).

Our results revealed that either under conditions of natural egress (after cpd1 egress arrest), or after treatment with pharmacological agents (zaprinast, saponin), two peaks of cytosolic Ca^2+^ increase occur in tachyzoites, the first one probably of intracellular store origin and the second peak associated with Ca^2+^ entry. In support of this statement, the second peak was absent after treatment with nifedipine, an agent previously found to be a Ca^2+^ entry inhibitor (28), and in *ΔPLP1* parasites, suggesting that this Ca^2+^ entry depends on the permeabilization of the host cell plasma membrane by the pore forming protein (PLP1). During natural egress, we consistently observed that, after an intracellular Ca^2+^ increase, a leading tachyzoite initiated egress via presumably secretion of PLP1, exit from the host cell, and finally rupture of the host plasma membrane. This was followed by the passive exit of the remaining tachyzoites. Our results showed a threshold for the Ca^2+^ increase that preceded egress, and we calculated this increase to be 300-500 nM [Ca^2+^]_i_ in the cytoplasm of tachyzoites. We stimulated host cell Ca^2+^ signaling with carbachol or histamine or used thapsigargin to cause Ca^2+^ increase in the cytosol of the host and parasite. These conditions, while causing an increase in [Ca^2+^]_i_ in tachyzoites, failed to reach the threshold needed for egress. These results demonstrated that parasites can replenish their intracellular stores while taking up Ca^2+^ from the host cell. Interestingly, treatment with thapsigargin lead to a Ca^2+^ increase in the host mitochondria surrounding the PV. Our experiments using patched host cells revealed that increasing Ca^2+^ concentrations in the host cell cytosol alone, presumably mimicking initial Ca^2+^ changes after plasma membrane rupture, were sufficient to stimulate egress, which was accelerated but not initiated by low K^+^ concentrations.

In our experimental set-up, stimulation of host Ca^2+^ signaling with known mammalian Ca^2+^ agonists resulted in host cytosolic Ca^2+^ increases of ∼1-2 μM. Interestingly, we saw that these low levels of Ca^2+^ are sufficient to stimulate Ca^2+^ influx into intracellular parasites followed by Ca^2+^ oscillations. Cytosolic Ca^2+^ oscillations in *T. gondii* tachyzoites were previously observed (6, 29), a fascinating phenomenon for which we do not have a molecular explanation. It is generally believed that Ca^2+^ oscillations arise from cyclical release and re-uptake of intracellularly stored Ca^2+^ (30), and a role for influx through plasma membrane channels has also been demonstrated to be important for the maintenance and delivery of Ca^2+^ into the cytosol. Because of the apparent digital nature of these Ca^2+^ oscillations (31), they would be perfectly suited for signaling specific biological responses such as secretion of micronemes, stimulation of motility and egress. It was shown that Ca^2+^ oscillations in extracellular tachyzoites loaded with Ca^2+^ dyes were associated with microneme discharge and bursts of motility (6). Changes in the Ca^2+^ oscillation patterns by pharmacological agents affected microneme secretion and motility (29, 32), highlighting the role Ca^2+^ oscillations in coordinating lytic cycle progression. In the case of intracellular tachyzoites, we believe that Ca^2+^ oscillations, while being insufficient to stimulate egress they serve to ensure the filling of intracellular stores. These oscillations are the result of Ca^2+^ release from the ER through an unknown channel, likely gated by IP_3_ (33), followed by re-uptake likely through the SERCA-Ca^2+^ ATPase (29) or the plasma membrane Ca^2+^ ATPase pump (34). The initial increase of cytosolic Ca^2+^ would be the result of uptake from the host through an unknown parasite plasma membrane channel and this initial Ca^2+^ increase would stimulate the activity of the PI-PLC at the plasma membrane (35, 36) to synthesize IP_3_, which would open the ER channel and release Ca^2+^ into the cytosol thereby potentiating the Ca^2+^ signals. However, cytosolic Ca^2+^ levels of the parasite are highly buffered (37), and they will be countered by mechanisms such as the PMCA-ATPase or the SERCA-Ca^2+^-ATPase to bring cytosolic Ca^2+^ back to normal basal levels (8). Another possible scenario would be that interactions between plasma membrane Ca^2+^ channels with ER receptors, plus Ca^2+^ itself (and other ions and signaling molecules) would result in PM Ca^2+^ channel openings, a phenomenon similar to what happens in muscle cells, where the Ryanodine Receptor (RyR) is directly or indirectly modulated by the L-type Ca^2+^ channel, CaV1.1/1.2, Ca^2+^ itself, other ions and small molecules, and proteins (38). Amplification of the initial Ca^2+^ signal occurs through activation of the RyR leading to Ca^2+^ release from the sarcoplasmic reticulum (5). This second scenario would require close proximity between the ER membrane and the plasma membrane because of the known poor diffusibility of Ca^2+^ in the cytoplasm of cells (39).

Within host cells parasites are stationary, non-motile, and surrounded by low Ca^2+^, yet activation by an intrinsic signal like phosphatidic acid, as recently proposed (40), would start a signaling cascade leading to a rise in cytosolic Ca^2+^ followed by stimulation of motility. Apicomplexan motility functions through a conserved, actomyosin motor that forms a complex termed the glideosome, the main molecular machinery involved in parasite motility, egress, and invasion (9). Within the glideosome, Regulatory Light Chains (MLC1), Essential Light Chains 1 and 2 (ELC1 and ELC2), Ca^2+^ binding proteins, interact either directly with Ca^2+^ or via Ca^2+^ dependent phosphorylation events to stimulate motility (41). The binding affinity of ELC is 37 μM (41), a concentration that would only be physiologically possible to reach if we accept/consider the presence of Ca^2+^ microdomains (confined regions of elevated Ca^2+^ at the site of activated/open channels) at the periphery of the parasite likely formed close to plasma membrane Ca^2+^ channels. This scenario supports the importance of Ca^2+^ influx for the stimulation of motility and egress, necessary to activate the glideosome while the overall cytosolic Ca^2+^ concentration would be maintained low.

It has been proposed that the high K^+^ content of the host cytosol blocks parasite egress (12). When the integrity of the host cell becomes compromised, the K^+^ concentration drops due to diffusion into the extracellular media. This drop in the concentration of K^+^ would induce a Ca^2+^ signal that would stimulate parasite egress through activation of a PI-PLC, though the molecules involved in this model are still unknown (12, 36). PI-PLCs of the delta type, like the type present in *T. gondii* (35) are highly sensitive to Ca^2+^ concentrations and are active at the Ca^2+^ levels present in the cytoplasm of cells (42). There is no direct connection between PI-PLC activity and K^+^ concentration but two genes in the *T. gondii* database (TGME49_238995 and TGME49_273380) are predicted to act as Ca^2+^-activated K^+^ channels. During intracellular growth, these channels could activate/open in response to Ca^2+^ release from intracellular stores. Initially, no conductance would occur because the potassium concentration of both parasite and host cytosol would be similar. Lysis of the host cell would result in decrease of the concentration of host cell and PV luminal K^+^, generating a gradient, and conductance of the ion with loss of K^+^ from the parasite, leading to an unbalance in the intracellular concentration of parasite intracellular K^+^ that would be counteracted by a plasma membrane mechanism such as a potassium/proton exchanger that would exchange K^+^ for H^+^. This mechanism would lead to acidification of the PV, which has been shown to precede egress (43). *T. gondii* expresses four predicted sodium proton exchangers and one of them was localized to the plasma membrane of *T. gondii (TgNHE1)* (TGME49_259200) (44), and a second one (TGME49_263480) is predicted to be at the plasma membrane according to ToxoDB LOPIT (45). It is possible that one of these exchangers could use K^+^ instead of Na^+^ and be responsible for the exchange activity. According to this potential scenario, Ca^2+^ would be the main trigger and K^+^ would have a modulatory role as we observed with the patch egress experiments.

In summary, we propose that as parasites grow and replicate, host Ca^2+^ derived from Ca^2+^ signaling is taken up by parasites through their plasma membrane and pumped into intracellular stores in order to keep them filled during repetitive rounds of replication (Fig. 7). Egress initiates via an unknown signal that induces release of Ca^2+^ from intracellular stores. Within the PV a leader parasite would respond first with the initial largest Ca^2+^ increase, which would reach the threshold needed for motility induction and secretion of microneme proteins. Release of PLP1 among other microneme proteins would contribute to the partial breakdown of the host cell (11). This allows for extracellular Ca^2+^ influx and a drop of host K^+^, two factors that would both contribute to egress of the remaining parasites. The cycle would continue in the remaining parasites: a Ca^2+^ spike would occur to reach the threshold for secretion of microneme proteins, complete breakdown of the PV membrane and host cell, and a second and more pronounced peak in Ca^2+^, culminating in egress of the entire PV of parasites. We found that the decrease in the concentration of K^+^ plays a role in timing parasite egress, and Ca^2+^ influx from the host cell or the extracellular milieu would be essential for parasite egress. The data from this work contributes to our knowledge of host-parasite interaction of *T. gondii* and aids in our understanding of the signaling responses that govern parasite egress. The use of whole-cell patching presents a novel methodology to study the role of ions involved in parasite egress, an essential component of the lytic cycle.

**Figure 7:**
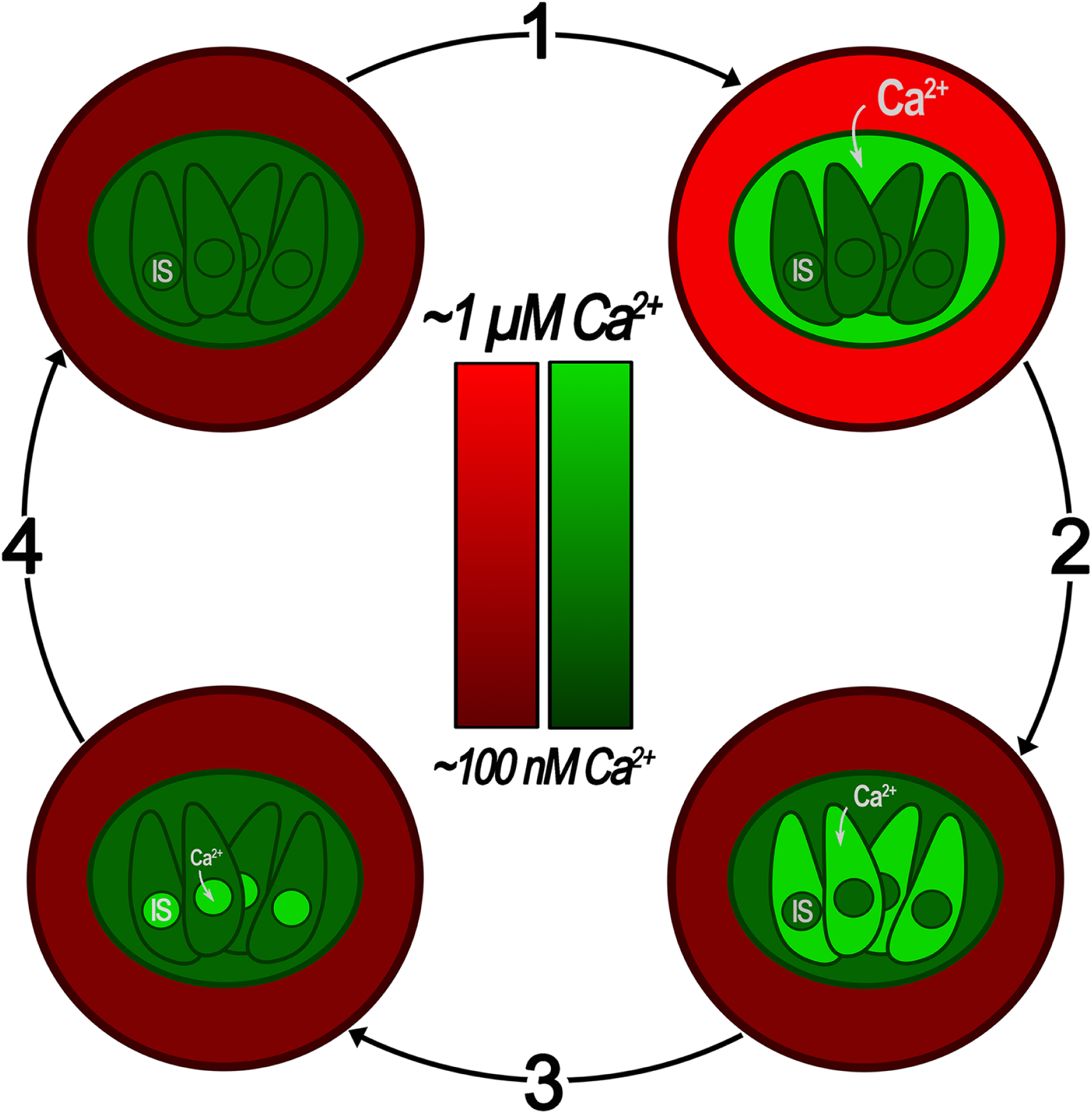
Model of Host Ca^2+^ Influx During Intracellular Growth. (1) A host cell Ca^2+^ signaling event triggers an increase in host cell cytosolic Ca^2+^. Given that the PV is in equilibrium with the host cell cytosol, PV Ca^2+^ rises simultaneously. (2) The rise in PV Ca^2+^ precedes Ca^2+^ influx into the parasite via an unknown channel thus causing a rise in the cytosolic Ca^2+^ of the parasite. (3) The increased cytosolic Ca^2+^ load of the parasite eventually returns back to basal levels as Ca^2+^ is pumped into the intracellular stores (IS). (4) The parasite continues replicating within the host cell while utilizing the Ca^2+^ influx from the host cell to maintain IS Ca^2+^ levels sufficient enough to be eventually utilized and released during egress.

## Materials and Methods

### Cell Culture

*T. gondii* tachyzoites (RH strain) were maintained in hTERT human fibroblasts (BD Biosciences) using Dulbeco’s modified essential media (DMEM) with 1% fetal bovine serum (FBS), as described previously (46). GCaMP6f-transfected tachyzoites were maintained under similar conditions, in the presence of 20 μM chloramphenicol. The selection-less strain of GCaMP6f was maintained without drug pressure. hTERT cells were maintained in high glucose DMEM with 10% fetal calf serum (FCS). HeLa cells (ATCC) were used for egress and whole-cell patch experiments and were maintained in DMEM supplemented with 10% FBS, 1 mM sodium pyruvate, and 2 mM L-glutamine. Cell cultures were grown at 37°C with 5% CO_2_. Parasites were purified by centrifugation and filtration through a Whatman 8 µM nuclepore membrane (GE Healthcare) followed by a second filtration step through a 5 µM nuclepore membrane. Filtered parasites were counted and centrifuged following the protocols for each experiment.

### Chemicals and Reagents

Transient transfections were performed using PolyJet that was purchased from SignaGen (http://signagen.com/). Plasmids for GCaMP6f (fast version of GCaMP6), R-GECO1.2, jGCaMP7f, and LAR-GECO1.2 were obtained from Addgene, and the plasmids were used for transient transfection in HeLa cells. The respective genes were cloned into the *T. gondii* expression vector pCTH3 and pDHFRTubGFP for chloramphenicol and selection-less stable expression of GCaMP6f in tachyzoites, respectively. Thapsigargin, ionomycin, saponin, dithiothreitol (DTT), histamine, Zaprinast, and all other chemicals were obtained from Sigma.

### Preparation of GECI-expressing Tachyzoites and HeLa Cells

GCaMP6f expressing parasites were obtained as described previously (7). Briefly, plasmids for expressing GCaMP6f in *T. gondii* were a gift from Kevin Brown and David Sibley. The coding DNA sequence for GCaMP6f was amplified by PCR and cloned into a *T. gondii* vector for expression downstream the tubulin promoter (pCTH3 and pDTGCaMP6f), using the BglII and AvrII restriction sites and adding a stop codon in front of the GFP sequence. The primers used were forward 5′-AGGCGTGTACGGTGGGAGGTC-3′ and reverse 5′CTTCCTAGGTTACTTCGCTGTCATCATTTG-3′ (Table S1). The Plasmids were electroporated into the RH strain parasites and clones were selected by chloramphenicol resistance. We then isolated cells with low fluorescence by cell sorting to eliminate those highly fluorescent cells in which the GCaMP6f could be buffering Ca^2+^ and preventing visualization of physiological changes in Ca^2+^ levels (47).

HeLa cells (5 x 10^5^) were grown on coverslips in high-glucose DMEM with 10% FBS. After 24 hours, cells were transfected with 1 μg of plasmid DNA encoding RGECO or LAR-GECO1.2 using PolyJet following the instructions of the manufacturer. 6-8 hours later, HeLa cells were infected with 1 x 10^6^ tachyzoites expressing GCaMP6f and were grown for 15-20 hours. Rosettes containing 4-8 parasites were used in all experiments. To construct a cell line of Hela cells stabling expressing the sensitive red GECI, jRGECO1a, the coding sequence of jRGECO1a (14) was PCR amplified with primers Fwd 5’-ACCGGTATGCTGCAGAACGAGCTTGCTCTTA -3’ and Rev 5’ -GAATTCGCCTACTTCGCTGTCATCATTTGTACA-3’ and cloned into the TOPO vector sequence. The resulting product was digested with AgeI and EcoRI restriction enzymes and cloned into the 2^nd^ generation lentivirus expression plasmid pUltra (Table S1). Transfection and cloning of a stable cell line of jRGECO1a was performed using previously established protocols (48). Briefly, 2^nd^ generation viral particles were produced in HEK293T cells, and the viral supernatant was overlaid on Hela cells for spinfection for 2 hr. Stable cell lines were enriched and selected using FAC’s sorting.

### Cytosolic Ca^2+^ Measurements

Extracellular tachyzoites of the RH strain were loaded with fura-2-AM as described previously (28). The parasites were washed twice in Ringer buffer (155 mM NaCl, 3 mM KCL, 2 mM CaCl_2_, 1 mM MgCl_2_, 3 mM NaH_2_PO_4_, 10 mM Hepes, pH 7.3, and 5 mM glucose), resuspended in the same buffer to a final density of 1 x 10^9^ cells/mL, and kept on ice. For fluorescence measurements, 50 µL-portions of the cell suspension were diluted in 2.5 mL of Ringer (2 x 10^7^ cells/mL final density) in a cuvette placed in a thermostatically controlled Hitachi 4500 spectrofluorometer. Excitation was at 485 nm and emission at 520 nm. Traces shown are representative of three independent experiments conducted on separate cell preparations. Calcium-defined conditions were determined by using EGTA or 1,2-bis(2-aminophenoxy)ethane-*N,N,N*′,*N*′-tetraacetic acid (BAPTA) and calcium chloride to reach specific concentrations of free calcium. Calcium-EGTA combinations were determined using Maxchelator software (https://somapp.ucdmc.ucdavis.edu/pharmacology/bers/maxchelator/downloads.htm).

### Egress Assays

Egress assays were done as described previously (47). HeLa cells were grown in high glucose DMEM with 10% FBS in 35 mm glass bottom dishes (MatTek) until confluency. 8-12 hours after transfection, HeLa cells were infected with 1 x 10^6^ GCaMP6f-expressing tachyzoites, replacing the media with high glucose DMEM with 1% FBS. Thirty hours after infection, parasitophorous vacuoles containing 4-8 parasites were observed by microscopy after washing them in the specific buffer for each experiment. Drugs were added 1 minute after the start of imaging in that same buffer at the concentrations indicated: ionomycin (0.005-1 μM), histamine (100 μM), thapsigargin (1 μM), saponin (0.01%), nifedipine (10 μM), and zaprinast (100 μM). Ringer buffer was used as extracellular buffer (EB). CaCl_2_ was omitted in the absence of extracellular calcium, and the media was supplemented with either 100 μM EGTA, 1 mM EGTA or 1 mM BAPTA. The composition of intracellular buffer (IB) is: 140 mM potassium gluconate, 10 mM NaCl, 2.7 mM MgSO_4_, 2 mM ATP (sodium salt), 1 mM glucose, 200 μM EGTA, 65 μM CaCl_2_ (90 nM free Ca^2+^), and 10 mM Tris/Hepes, pH 7.3. The parasites were imaged at 37°C. Fluorescence images were captured using an Olympus IX-71 inverted fluorescence microscope with a Photometrix CoolSnapHQ charge-coupled device (CCD) camera driven by DeltaVision software (Applied Precision). Images were collected using time-lapse mode with an acquisition rate of at least 2-3 seconds during 10-20 minutes. Images were converted into videos using SoftWorx suite 2.0 software from Applied Precision. Fiji was used for the analysis of the video data. Fluorescence tracings were produced by drawing an ROI around the host or parasite of interest and measuring the mean gray fluorescence. Prism was used for statistical analysis.

For natural egress, 3.5 x 10^5^ Hela cells stabling expressing the red GECI jRGECO1a (14) were plated on 35 mm glass bottom MatTek dishes in DMEM-HG with 10% FBS. 24 h later cells were infected with 2.75 x 10^6^ parasites, and on day three the parasites were synchronized using 1 μM compound, and one dish was treated with 2 μL of DMSO as a vehicle control. On day four lysis of the control dish was examined for 70-80% lysis, and the synchronized dishes were used for video microscopy. The dishes were washed once with pre-warmed Ringer buffer, then again with pre-warmed Ringer buffer for 1 minute and finally a solution of Ringer buffer supplemented with 2 mM CaCl_2_ was added before commencing imaging.

### Whole cell patch

Whole cell patch recording was applied according to the method previously described (49, 50). Briefly, coverslips with infected HeLa cells were transferred into a perfusion chamber (RC-26GLP, Warner Instruments, USA) on a stage of an inverted IX51 Olympus microscope. Growth media was immediately replaced with an extracellular buffer prior to the start of the recordings, and cells were kept at room temperature for no more than 2 hours. Coverslips were replaced every 2 hours. Intracellular thin-walled recording capillars (1.5-mm outer diameter, 1.17-mm inner diameter) made of borosilicate glass (World Precision Instruments, Sarasota, FL) were used to pull patch pipettes (3–4 MΩ) on a horizontal Flaming/Brown micropipette puller (P-97, Sutter Instruments) and then were backfilled with intracellular solution. Recordings from HeLa cells were obtained with an Axopatch 200B amplifier (Molecular Devices) using high-resolution videomicroscopy (Ameriscope). The voltage output was digitized at 16-bit resolution, 20 kHz, and filtered at 1 kHz (Digidata 1550A, Molecular Devices). All experiments were performed at −60 mV holding membrane potential (Vh) which is close to values reported as physiological for HeLa cells (51) to prevent activation of voltage-gated calcium channels. After establishing a “seal” (>1 MΩ) slight negative pressure was applied together with “ZAP” electrical pulse (0.5-5 ms, +1.3V) to break cell membrane. The time from membrane opening (start of pipette/cytosol solution exchange) to egress was visually recorded. All statistical analyses were conducted using GraphPad Prism 5 (GraphPad Software, San Diego, CA). All values are expressed as means ± SE. Between-group differences were compared using unpaired *t*-tests. Differences were considered statistically significant at *P* < 0.05, and *n* refers to the number of cells.

HeLa cells grown in high glucose DMEM with 10% FBS on 22 x 40 mm glass coverslips until 70% confluency were infected with 1 x 10^6^ GCaMP6f-expressing tachyzoites. 24 h post infection coverslips were placed in an electrophysiology recording chamber (∼1 ml) and bathed in extracellular solution (mM): 140 NaCl, 5 KCl, 1 MgSO_4_, 1.8 CaCl_2_, 10 Hepes, pH 7.5 adjusted with NaOH/HCl. Composition of “high potassium” intracellular pipette solution (mM): 140 KCl, 5 NaCl, 2 MgCl_2_, 1 Glucose, 10 Hepes, 1 BAPTA, pH 7.3 adjusted with KOH/HCl. “Low potassium” solution was prepared as described above with equimolar substitution of 130 mM KCl with choline chloride (ChCl). Disodium ATP (2 mM) was added to all intracellular solutions to maintain cell energy supply. Solutions with different free Ca^2+^ concentrations were prepared by adding of Ca^2+^ and BAPTA at proportions calculated with Webmaxc software (Stanford University, USA). For control experiments in order to exclude the role of host cell extracellular calcium entry that may contribute to *T. gondii* egress, we used modified extracellular solution with MgCl_2_ and CaCl_2_ adjusted to 5 mM and 0.1 mM, respectively.

### Quantification and Statistical Analysis

Statistical analysis of fluorescence images was performed using FIJI/ImageJ (52). Briefly, images were background subtracted and normalized using an average of the first 5 frames of imaging. ΔF (F_max_/F_min_) represents the highest fold change over baseline in fluorescence after addition of stimuli or reagent. Figures were constructed using Prism analysis suite and error bars represent the standard error of the mean S.E.M. of three independent experiments. Significant differences were only considered if P values were < 0.05, where *p < 0.05; **p < 0.01; ***p < 0.001; and ****p < 0.0001. NS designates when the comparison is not statistically significant. Experiment-specific statistical information is provided in the figure legends or associated method details including trials (n), standard error of the mean SEM, and statistical test performed.

## Supplemental Material

**Fig S1. Impact of Host Histamine Signaling on *T. gondii***: A) Hela cells transiently expressing the red GECI RGECO were infected with *cyto*-GCaMP6f expressing parasites.; B) Stable Hela cell lines expressing jRGECO1a were infected with PV localized P30-jGCaMP7f expressing parasites; C) Hela cells infected with P30-RGECO *cyto*-GCaMP6f parasites. Under all conditions 100 μM Histamine was added at about 1 min mark and the response was recorded. Tracings to the right of each panel series show fluorescence fluctuations of single parasites (*green*) or of a delineated region of interest in the host cells (*red*) or PV (green) from 3 independent trials for each condition. Bar graphs represent quantification of the average ΔF values of three independent trials (Red, jRGECO1a or RGECO, *right axis*) (Green, GCaMP6f or jGCaMP7f, *left axis*). Dashed white outlines indicate the area used as a region of interest to analyze the fluorescence tracings. Dashed white outlines highlight GCaMP6f-expressing parasites whose fluorescence tracings were used for analysis. Dashed Red outlines indicate the region of the host cell that was used to analyze for the RGECO or jRGECO1a channel. Numbers at the upper right of each panel indicate the time frame of the video.

**Fig S2. ΔPLP1 Mutants Lack the Second Ca**^**2+**^ **Peak During Egress:** A) Egress was induced in Hela cells infected with ΔPLP1 GCaMP6f parasites using 100 μM Zaprinast; B & C) Representative fluorescence tracings of ΔPLP1 GCaMP6f after addition of 100 µM Zaprinast for egress and non-egressing parasites respectively. Notice, two peaks are visible for the parasites that were successfully able to egress and only one peak was visible for the parasites that were unable to egress.

**Fig S3. Calibration of Ca**^**2+**^ **Threshold:** (A) RH tachyzoites (5 x 10^7^) were loaded with FURA2-AM and resuspended in 2.5 mL of Ringer’s Buffer supplemented with 100 μM EGTA. The cytosolic Ca^2+^ response to the addition of various concentrations of ionomycin is shown. The figure shows the average of at least 2 trials of each condition; (B) Quantification of Ionomycin Titration Curve. The first 15 seconds of the slope after addition of ionomycin was determined and averaged for at least two independent trials.

**Video S1. Carbachol Addition in GCaMP6f Infected Host Cells:** Hela cells transiently expressing cytosolic localized RGECO were infected with cytosolic GCaMPf6 expressing parasites. 1 mM Carbachol was added at approximately the 1 min mark. (Fig. 1B shows still images from this video).

**Video S2. Thapsigargin Stimulation in LAR-GECO Expressing Host Cells:** Hela cells transiently expressing the low affinity mitochondrial Ca^2+^ indicator LAR-GECO1.2 were infected with cytosolic GCaMP6f expressing parasite. 2 μM Thapsigargin was added at approximately the 1 min mark. (Fig. 2B shows still images from this video).

**Video S3. Two-peaks of Ca**^**2+**^ **contribute to Parasite Egress:** Hela cells infected with cytosolic GCaMP6f parasites were treated with 100 μM of Zaprinast in media supplemented with 2 mM Ca^2+^. (Fig. 3E shows still images from this video).

**Video S4. Natural Egress in Cpd 1 Synchronized Parasites:** Hela cells stabling expressing cytosolic jRGECO1a infected with cytosolic GCaMP6f expressing parasites. After 24 hrs post infection 1 μM cpd1 was added. 24 hr post cpd1 addition parasite egress was imaged after cpd1 washout. (Fig. 4A shows still images from this video).

**Video S5. Whole-Cell Patch under High K**^**+**^ **and High Ca**^**2+**^ **Conditions:** Hela cells infected with GCaMP6f parasites were whole-cell patched with a patch pipette solution comprising 140 mM K^+^ and 10 μM Ca^2+^. Note parasite egress is slow, methodical, and occurring one-by-one in comparison to detergents and ionophores. Video presented in real time. (Fig. 6B shows still images from this video).

**Video S6. Whole-Cell Patch under Low K**^**+**^ **and High Ca**^**2+**^ **conditions:** Hela cells infected with GCaMP6f parasites were whole-cell patched with a patch pipette solution comprising 10 mM K^+^, 130 mM choline chloride, and 10 μM Ca^2+^. Again, parasites egress slower in comparison to detergents and ionophores, but more rapidly in comparison to high K^+^ conditions. Parasite egress occurred in which all parasites egressed at once under these conditions. Video presented in real time. (Fig. 6C shows still images from this video).

**Table S1. Plasmids used in this work**

## Acknowledgements

We would like to thank Alex W. Chan and Dr. Sebastian Lourido for the protocol of natural egress using compound 1; Dr. Vern Carruthers for the ΔPLP1 mutant parasites; Dr. Diego Huet for reading the manuscript; Daniel Williamson for assisting on the egress of the ΔPLP1 mutant; Beejan Asady for technical assistance; Julie Nelson for the FAC’s sorting assistance; Dr. Kandasamy for technical assistance on the use of the microscopes at the Biomedical Microscopy Core. This work was supported by an NIH grant R01AI128356 to SNJM. SAV and EP were partially supported by fellowships (pre-doc and post-doc respectively) through a Training Grant in Tropical and Emerging Global Diseases (T32AI060546).

## Authors Contributions

Conceived and designed the experiments: SAV, CAM, SNJM. Performed the experiments: SAV, CAM, ZHL, MAHT and EP. Analyzed the data: SAV, CAM and SNJM. Contributed reagents/materials/analysis tools: SAV, CAM, ZHL and SNJM. Wrote the paper: SAV and SNJM.

