## Supplemental figure legends for "The Role of Potassium and Host Calcium Signaling in *Toxoplasma gondii* egress"

Stephen A. Vella et al

#### Supporting Figures Legends

**Fig. S1. Impact of Host Histamine Signaling on *T. gondii*:** A) Hela cells transiently expressing the red GECI RGECO were infected with *cyto*-GCaMP6f expressing parasites.; B) Stable Hela cell lines expressing jRGECO1a were infected with PV localized P30-jGCaMP7f expressing parasites; C) Hela cells infected with P30-RGECO *cyto*-GCaMP6f parasites. Under all conditions 100 μM Histamine was added at about 1 min mark and the response was recorded. Tracings to the right of each panel series show fluorescence fluctuations of single parasites (*green*) or of a delineated region of interest in the host cells (*red*) or PV (green) from 3 independent trials for each condition. Bar graphs represent quantification of the average ΔF values of three independent trials (Red, jRGECO1a or RGECO, *right axis*) (Green, GCaMP6f or jGCaMP7f, *left axis*). Arrowheads indicate the area used as a region of interest to analyze the fluorescence tracings. White arrows highlight GCaMP6f-expressing parasites whose fluorescence tracings were used for analysis. Red arrows indicate the region of the host cell that was used to analyze for the RGECO or jRGECO1a channel. Numbers at the upper right of each panel indicate the time frame of the video.

**Fig. S2. ΔPLP1 Mutants Lack the Second $Ca^{2+}$ Peak During Egress:** A) Egress was induced in Hela cells infected with ΔPLP1 GCaMP6f parasites using 100 μM Zaprinast; B & C) Representative fluorescence tracings of ΔPLP1 GCaMP6f after addition of 100 μM Zaprinast for egress and non-egressing parasites respectively. Notice, two peaks are visible for the parasites that were successfully able to egress and only one peak was visible for the parasites that were unable to egress. D) Natural Egress of ΔPLP1 GCaMP6f parasites. ΔPLP1 GCaMP6f parasites were synchronized with 1 μM cpd1 for 24 hrs to inhibit egress. Post washout parasite egress can be visualized microscopically E) Fluorescence tracings of ΔPLP1 GCaMP6f parasites after cpd1 washout. Note a second peak does not occur as parasites are defective in egressing, and the jRGECO1a signal does not change because the host cell remains intact.

**Fig. S3. Calibration of $Ca^{2+}$ Threshold:** (A) RH tachyzoites (5 x $10^7$) were loaded with FURA2-AM and resuspended in 2.5 mL of Ringer's Buffer supplemented with 100 μM EGTA. The cytosolic $Ca^{2+}$ response to the addition of various concentrations of ionomycin is shown. The figure shows the average of at least 2 trials of each condition; (B) Quantification of Ionomycin Titration Curve. The first 15 seconds of the slope after addition of ionomycin was determined and averaged for at least two independent trials.

### Figure S1

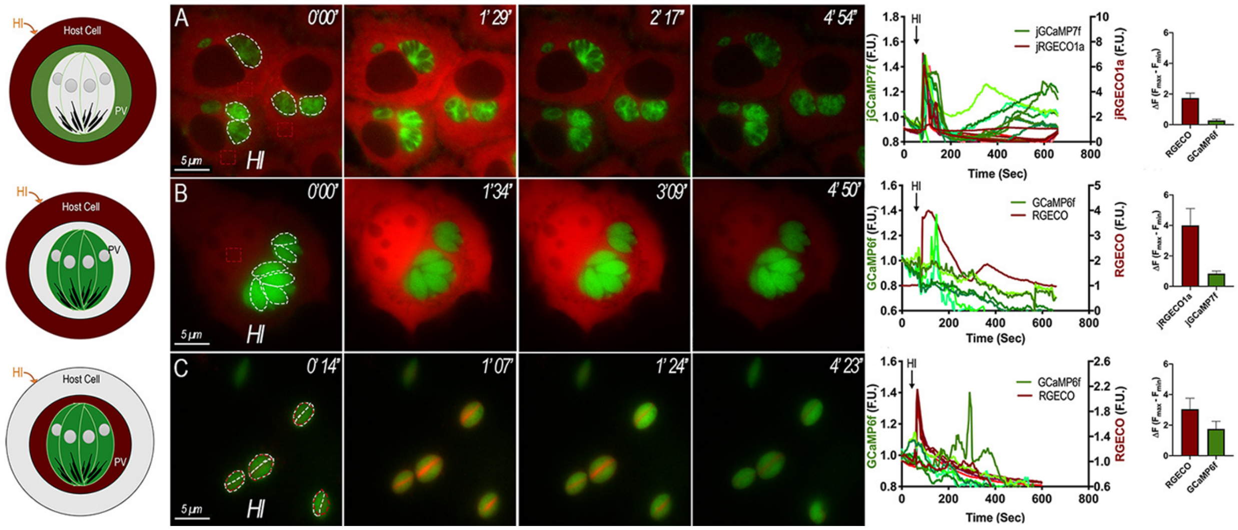

HI
Host Cell
PV
A
0'00"
1' 29"
2' 17"
4' 54"
5 µm
HI
jGCaMP7f
jRGECO1a
jGCaMP7f (F.U.)
jRGECO1a (F.U.)
Time (Sec)
RGECO
GCaMP6f
B
0'00"
1'34"
3'09"
4' 50"
GCaMP6f
RGECO
GCaMP6f (F.U.)
RGECO (F.U.)
jRGECO1a
jGCaMP7f
C
0' 14"
1' 07"
1' 24"
4' 23"

#### Figure S2

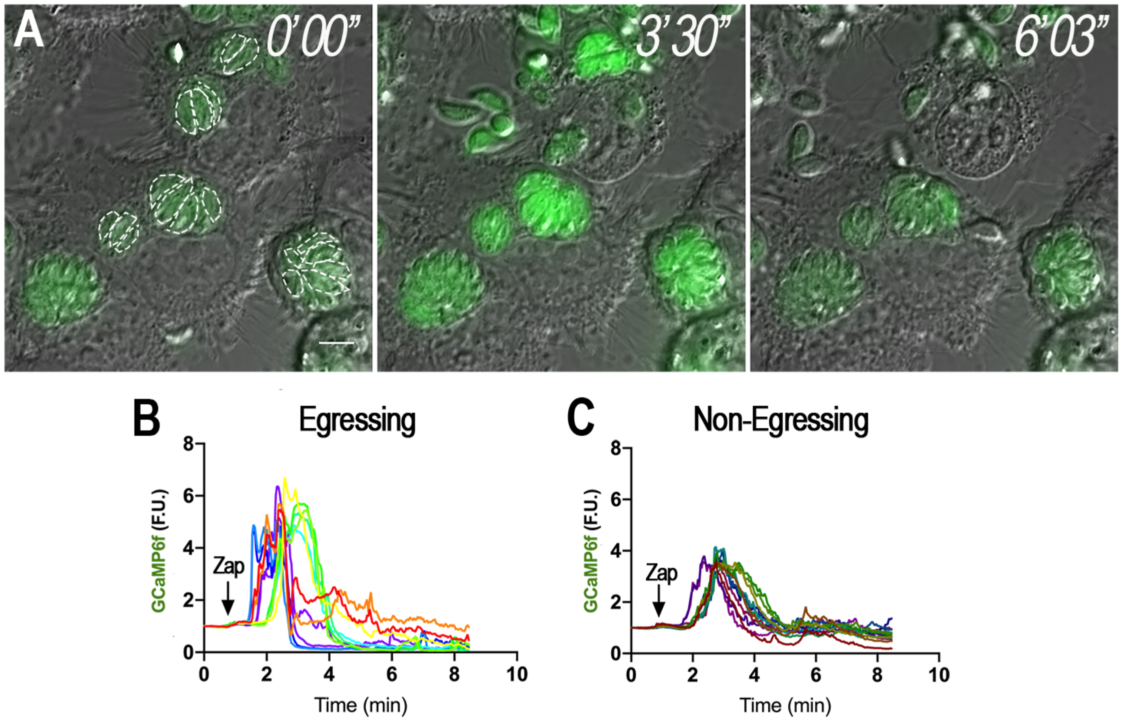
A
0'00"
3'30"
6'03"
B
Egressing
GCaMP6f (F.U.)
Zap
Time (min)
C
Non-Egressing
GCaMP6f (F.U.)
Zap
Time (min)

#### Figure S3

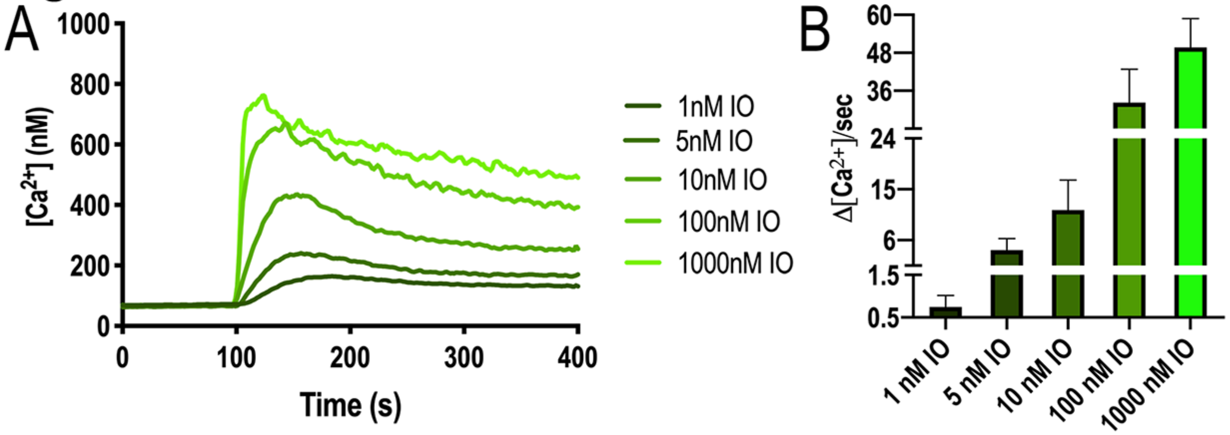
A
1000
800
600
400
200
0
[Ca2+] (nM)
0
100
200
300
400
Time (s)
1nM IO
5nM IO
10nM IO
100nM IO
1000nM IO
B
60
48
36
24
15
6
1.5
0.5
Δ[Ca2+]/sec
1 nM IO
5 nM IO
10 nM IO
100 nM IO
1000 nM IO

#### Supplemental Videos Legends

**Video S1. Carbachol Addition in GCaMP6f Infected Host Cells:** Hela cells transiently expressing cytosolic localized RGECO were infected with cytosolic GCaMPf6 expressing parasites. 1 mM Carbachol was added at approximately the 1 min mark. (Fig. 1B shows still images from this video).

**Video S2. Thapsigargin Stimulation in LAR-GECO Expressing Host Cells:** Hela cells transiently expressing the low affinity mitochondrial $Ca^{2+}$ indicator LAR-GECO1.2 were infected with cytosolic GCaMP6f expressing parasite. 2 μM Thapsigargin was added at approximately the 1 min mark. (Fig. 2B shows still images from this video).

**Video S3. Two-peaks of $Ca^{2+}$ contribute to Parasite Egress:** Hela cells infected with cytosolic GCaMP6f parasites were treated with 100 μM of Zaprinast in media supplemented with 2 mM $Ca^{2+}$. (Fig. 3E shows still images from this video).

**Video S4. Natural Egress in Cpd 1 Synchronized Parasites:** Hela cells stabling expressing cytosolic jRGECO1a infected with cytosolic GCaMP6f expressing parasites. After 24 hrs post infection 1 μM cpd1 was added. 24 hr post cpd1 addition parasite egress was imaged after cpd1 washout. (Fig. 4A shows still images from this video).

**Video S5. Whole-Cell Patch under High $K^+$ and High $Ca^{2+}$ Conditions:** Hela cells infected with GCaMP6f parasites were whole-cell patch clamped with a patch pipette solution comprising 140 mM $K^+$ and 10 μM $Ca^{2+}$. Note parasite egress is slow, methodical, and occurring one-by-one in comparison to detergents and ionophores. Video presented in real time. (Fig. 6B shows still images from this video).

**Video S6. Whole-Cell Patch under Low $K^+$ and High $Ca^{2+}$ conditions:** Hela cells infected with GCaMP6f parasites were whole-cell patch clamped with a patch pipette solution comprising 10 mM $K^+$, 130 mM choline chloride, and 10 μM $Ca^{2+}$. Again, parasites egress slower in comparison to detergents and ionophores, but more rapidly in comparison to high $K^+$ conditions. Parasite egress occurred in which all parasites egressed at once under these conditions. Video presented in real time. (Fig. 6C shows still images from this video).

#### Table S1: Plasmids used in this work

| Strain, Plasmid, or oligonucleotide | Relevant Characteristics | Source or Reference |
|---|---|---|
| **Plasmid** | | |
| pGP-CMV-GCaMP6f | Transient expression of the GECI GCaM6f in mammalian cells | [1] |
| pCMV-NLS-R-GECO | Transient expression of the GECI R-GECO in mammalian cells | [2] |
| pGP-CMV-NES-jRGECO1a | Transient expression of the GECI jRGECO1a in mammalian cells | [3] |
| pGP-CMV-jGCaMP7f | Transient expression of the GECI jGCaMP7f in mammalian cells | [4] |
| pCMV-mito-LAR-GECO1.2 | Transient expression of the GECI LAR-GECO1.2 in the mitochondria of mammalian cells | [5] |
| ptub_IEα-DsRed_DHFR_sag1CATsag1 | Plasmid vector for expressing proteins in the lumen of the PV | [6] |
| pCTH3 | Random Integration of a chloramphenicol selection plasmid expressing 3xHA epitopes | [7] |
| ptub_IEα-R-GECO_DHFR_sag1CATsag1 | Expressing the GECI R-GECO in the lumen of the PV | [8] |
| pUltra | Lentivirus expression of a gene of interest | [9] |
| jRGECO1a-pUltra | Lentivirus expression of the Red GECI jRGECO1a | This study |
| **Oligonucleotides** | | |
| **To construct PV-jGCaMP7f** | | |
| SV229-jGCaMP7f-GIB-FWD | GGGCTGCAatgggttctcatcatcatc | This study |
| SV72-jRCaMP1b-Gib-R | ccgggacgtcgtacgggtacctaggCTTCGCTGTCATCATTTGTAC | This study |
| SV77-P30-Gib-Fwd | TTTCTTGAATTCCCTTTTAGATCTATGTTTCCGAAGGCAGTGAGACG | This study |
| SV-228-P30-Gib-Rev | gaacccatTGCAGCCCCGGCAAACTC | This study |
| SV225-pCTH3-Gib-FWD | cagcgaagCctaggtacccgtacgac | This study |
| SV226-Tub-Gib-Rev | CGGAAACATAGATCtaaaagggaattcaagaaaaaatg | This study |
| **To construct jRGECO1a-pUltra** | | |
| SV155V2-jRGECO1a-AgeI-Fwd | aCCGGTATGCTGCAGAACGAGCTTGCTCTTA | This study |

| | | |
|---|---|---|
| SV156-jRGECO1a-ECORI-Rev | GAATTCGCCTACTTCGCTGTCATCATTTG TACA | This study |
| **Strain** | | |
| RH GCaMP6f | Non-selectable drug RH strain expressing GCaMP6f | [8] |
| RH PV-jGCaMP7f | Expression of jGCaMP7f in the lumen of the PV | This study |
| RH PV-RGECO GCaMP6f | Expression of PV-RGECO in the lumen of the PV and cytosolic expression of GCaMP6f | [8] |
